# Comparison of biological features of wild European rabbit mesenchymal stem cells derived from different tissues

**DOI:** 10.1101/2021.12.13.472420

**Authors:** Alexandra Calle, María Zamora-Ceballos, Juan Bárcena, Esther Blanco, Miguel Ángel Ramírez

**Affiliations:** Departamento de Reproducción Animal, INIA, CSIC, 28040 Madrid, Spain; Instituto Centro de Investigación en Sanidad Animal (CISA), INIA, CSIC, 28130 Madrid, Spain

**Keywords:** endangered species, European rabbit, mesenchymal stem cells

## Abstract

Although the European rabbit is an “endangered” species and a notorious biological model, the analysis and comparative characterization of new tissue sources of rabbit mesenchymal stem cells (rMSCs) has not been well studied. Here we report for the first time the isolation and characterization of rMSCs derived from an animal belonging to a natural rabbit population within the species native region. New rMSC lines were isolated from different tissues: oral mucosa (rOM-MSC), dermal skin (rDS-MSC), subcutaneous adipose tissue (rSCA-MSC), ovarian adipose tissue (rOA-MSC), oviduct (rO-MSC), and mammary gland (rMG-MSC). The six rMSC lines showed plastic adhesion with fibroblast-like morphology and were all shown to be positive for CD44 and CD29 expression (characteristic markers of MSCs), and negative for CD34 or CD45 expression. In terms of pluripotency features, all rMSC lines expressed NANOG, OCT4, and SOX2. Furthermore, all rMSC lines cultured under osteogenic, chondrogenic, and adipogenic conditions showed differentiation capacity. In conclusion, this study describes the isolation and characterization of new rabbit cell lines from different tissue origins, with a clear mesenchymal pattern. We show that rMSC do not exhibit differences in terms of morphological features, expression of the cell surface, and intracellular markers of pluripotency and *in vitro* differentiation capacities, attributable to their tissue of origin.

## 1. Introduction

The European rabbit (*Oryctolagus cuniculus*) originated in the Iberian Peninsula. The species expanded through anthropogenic dispersal to Europe, Australia, New Zealand, the Americas, and North Africa [1]. A single origin of domestication led to lower levels of genetic diversity within domestic rabbits than those found in wild populations [2]. Thus, natural populations within the European rabbit native region, the Iberian Peninsula, are considered a reservoir of genetic diversity [3]. Over the last 70 years, the European rabbit populations, both in its native and non-native ranges, have suffered declining population trends as reflected in the successive declaration statuses by IUCN: 1996 - Lower Risk / least concern (LR / LC); 2008 - Near Threatened (NT) and finally 2018 - Endangered (https://www.iucnredlist.org/species/41291/170619657). The emergence of two viral diseases: myxomatosis and rabbit hemorrhagic disease (RHD) was the primary cause of the decline [4–7]. In this context, the European rabbit provides a suitable system to study host–viral pathogen interactions, coevolution, and ecological effects [8–11].

The European rabbit has been extensively studied as a laboratory model for human diseases in both biomedical and fundamental research [12]. After mice (60.9%) and rats (13.9%), the laboratory rabbit is the third most used mammal model in animal experimentation (3.12%) within the EU. Furthermore, the highest increase in comparison to the previous report in 2008 is noted for fish and rabbits (EU report 2013: https://eur-lex.europa.eu/legal-content/EN/TXT/PDF/?uri=CELEX:52013DC0859). However, our current knowledge of important aspects such as rabbit immunogenetics or the characterization of mesenchymal stem cells (MSCs) lags behind that of other relevant model species, like the mouse.

MSCs are widely utilized in therapy because of their immunomodulatory properties [13], but their potential application in infectious viral diseases is less explored. MSCs are susceptible to infection by a wide variety of RNA and DNA viruses both *in vitro* and *in vivo* [14–17] Despite their permissiveness to some viruses’ entry, some evidence has also emerged that MSCs can mitigate viral infection via the upregulation of antiviral mechanisms. MSC-released miRNAs have been reported to inhibit hepatitis C virus replication [18]. MSCs can interact with and influence both the innate and adaptive immune cells, potentially altering the outcome of the response to a viral infection. While there is already information in the literature regarding the use of MSCs and/or their secretome as a therapy in several animal models of viral diseases, their applications in rabbits have still not been assessed [19].

MSCs have been found in some rabbit tissues (adipose, gingival, bone marrow, Wharton’s jelly, amniotic fluid) exhibiting stem cell characteristics such as plastic adherence, multilineage differentiation capacity, MSC marker expression, and pluripotent gene expression [20–23]. Postnatal organs and tissues are good MSC sources; however, each source of MSCs has a different degree of differentiation potential and expression of a different set of stem cell-related markers, as well as other important characteristics such as high proliferation, immunomodulation, and allo- and xeno-transplantation ability.

Rabbit MSCs (rMSC) have been widely used as preclinical models for orthopedic, specifically bone, articular cartilage, ligament reconstruction, and spinal fusion, as well as cardiovascular regenerative medicine strategies [24–27]. Although rMSCs are being investigated in these models, they have not been fully characterized in terms of immunophenotype and differentiation potential, and many tissues have not yet been explored as a source of MSC in the rabbit. Moreover, all the previous rMSC reported in the literature had been obtained from domestic New Zealand White (NZW) line rabbits. Here we report for the first time the isolation and characterization of rMSC obtained from a female specimen belonging to a wild rabbit population of the species native range, the Iberian Peninsula.

The main goal of the present work was to perform the characterization of rMSC lines isolated from different tissues, in terms of morphological features, expression of mesenchymal and pluripotency-related markers, and *in vitro* chondrogenic, osteogenic and adipogenic differentiation capacities.

## 2. Results

### 2.1. Morphological features of different rMSC

As shown in Figure 1, we could successfully isolate rMSC from ovarian adipose tissue, subcutaneous adipose tissue, dermal skin, oral mucosa, oviduct, and mammary gland of an adult female wild rabbit.

**Figure 1.**
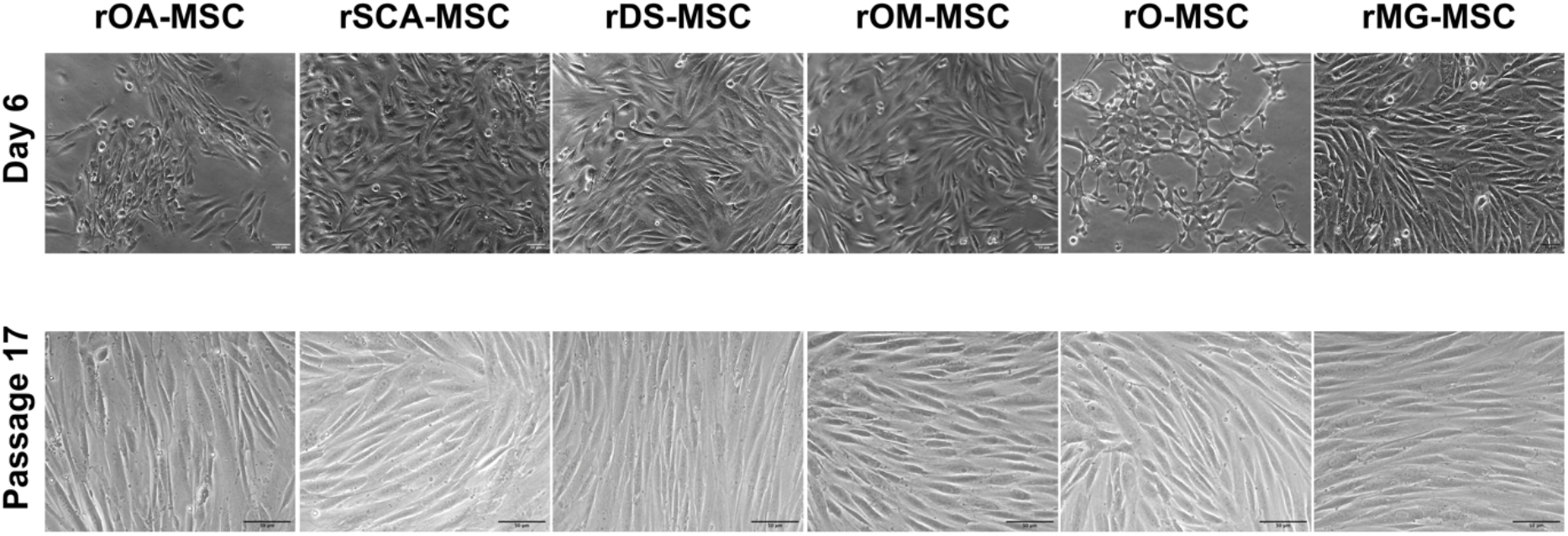
Phase-contrast images of different rabbit rMSC lines at 6 days of culture of Passage 0 (upper panels; ×100 magnification) and Passage 17 (lower panels; x200 magnification). Bars = 50 µm.

In primary culture, rMSCs of all six sources adhered to the plastic surface of culture dishes, exhibiting a mixture of round, spindle, or elongated shape morphologies (Figure 1-upper panels). However, after the first cell passage, cells formed a homogeneous population of fibroblast-like adherent cells (Figure 1—lower panels).

### 2.2. Immunophenotypic characterization by flow cytometry

For further characterization of all six types of rMSCs, some characteristic cell surface markers were assessed by flow cytometry (Figure 2). Cell surface CD44, a characteristic marker of rMSCs, was expressed at detectable levels in all cell lines. In none of the rMSC lines, expression of hematopoietic lineage markers like CD34 and CD45 was detected.

**Figure 2.**
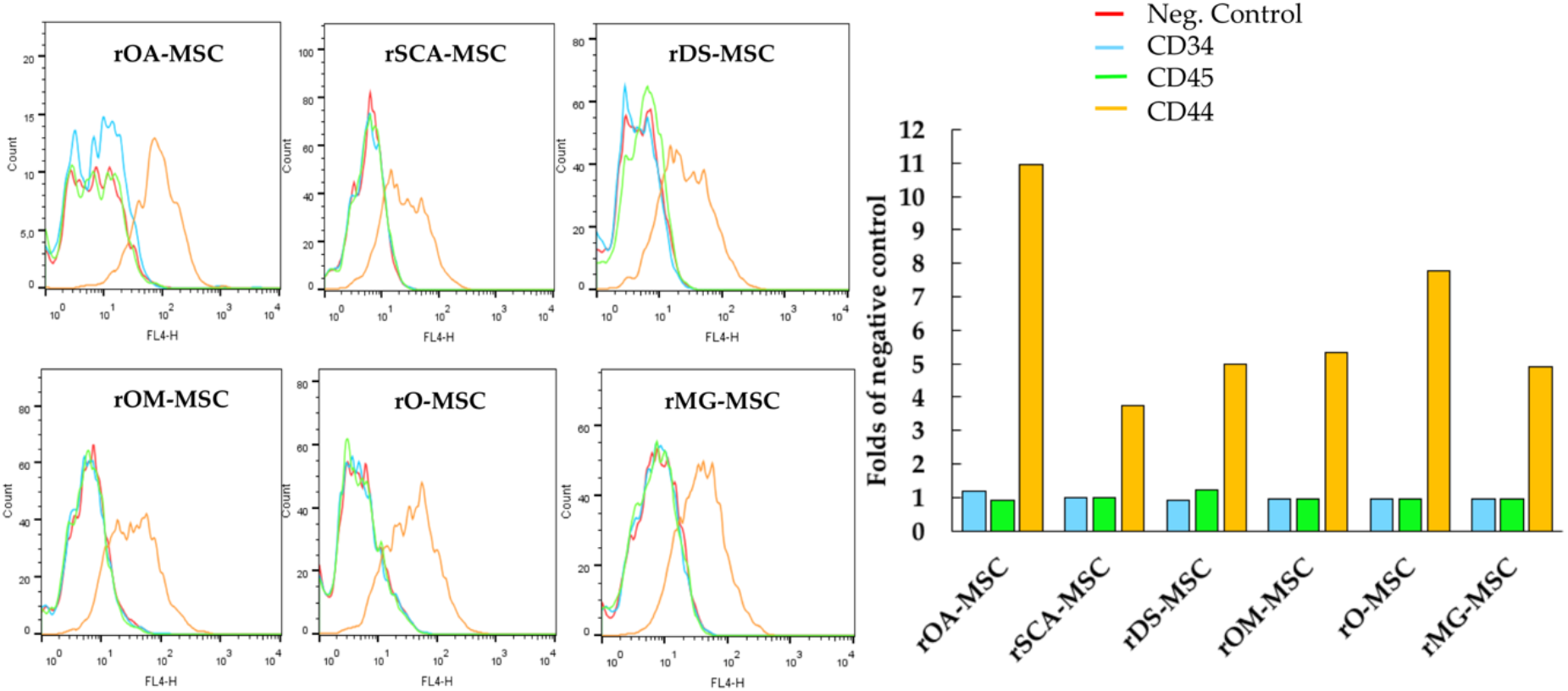
Analysis by flow cytometry of the expression levels of cell surface markers CD34, CD44, and CD45 in rOA-MSC, rSCA-MSC, rDS-MSC, rOM-MSC, rO-MSC, rMG-MSC, and RK13 rabbit kidney epithelial cells. Plots depict the log of fluorescence intensity for each sample and bar charts the mean fluorescence intensity related to the negative control sample.

### 2.3. Gene expression of surface and pluripotency markers detected by RT-PCR

RT-PCR analyses confirmed the expression in all rMSCs lines of those CD surface markers that are characteristic of MSCs (CD29 and CD44) (Figure 3). In addition, CD34 and CD45, hematopoietic lineage markers, could not be detected at the mRNA level in the tested samples. Pluripotent markers OCT4, SOX2, and NANOG were detected by RT-PCR to be expressed in all rMSC lines.

**Figure 3.**
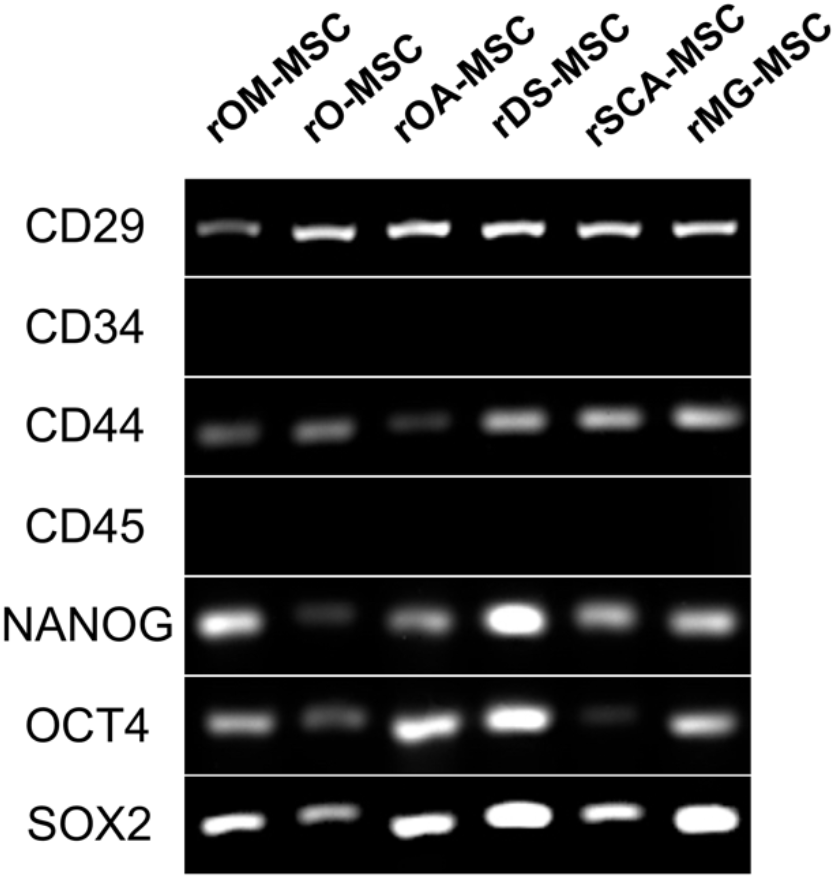
RT-PCR analysis of surface and pluripotency marker gene expression in the different rMSC. A panel of surface markers including CD29, CD34, CD44, and CD45 was used for the identification of rMSC. Pluripotency genes expression (NANOG, OCT4, and SOX2) was detected in the different rMSC.

### 2.4. In vitro differentiation of rMSCs

As shown in Figure 4, all rMSC lines cultured under chondrogenic conditions showed the presence of acidic proteoglycan synthesized by chondrocytes that were demonstrated at the monolayer of cells by Alcian blue staining, which appeared in most cases as stained nodules typical from cartilaginous tissue. Cells cultured under osteogenic conditions presented remarkable calcium deposits, indicating a high osteogenic differentiation potential of these lines. All rMSC lines obtained were also able to differentiate to adipocytes when cultured under adipogenic conditions, presenting the formation of cytoplasmic lipid droplets visualized after the Oil red O solution stain.

**Figure 4.**
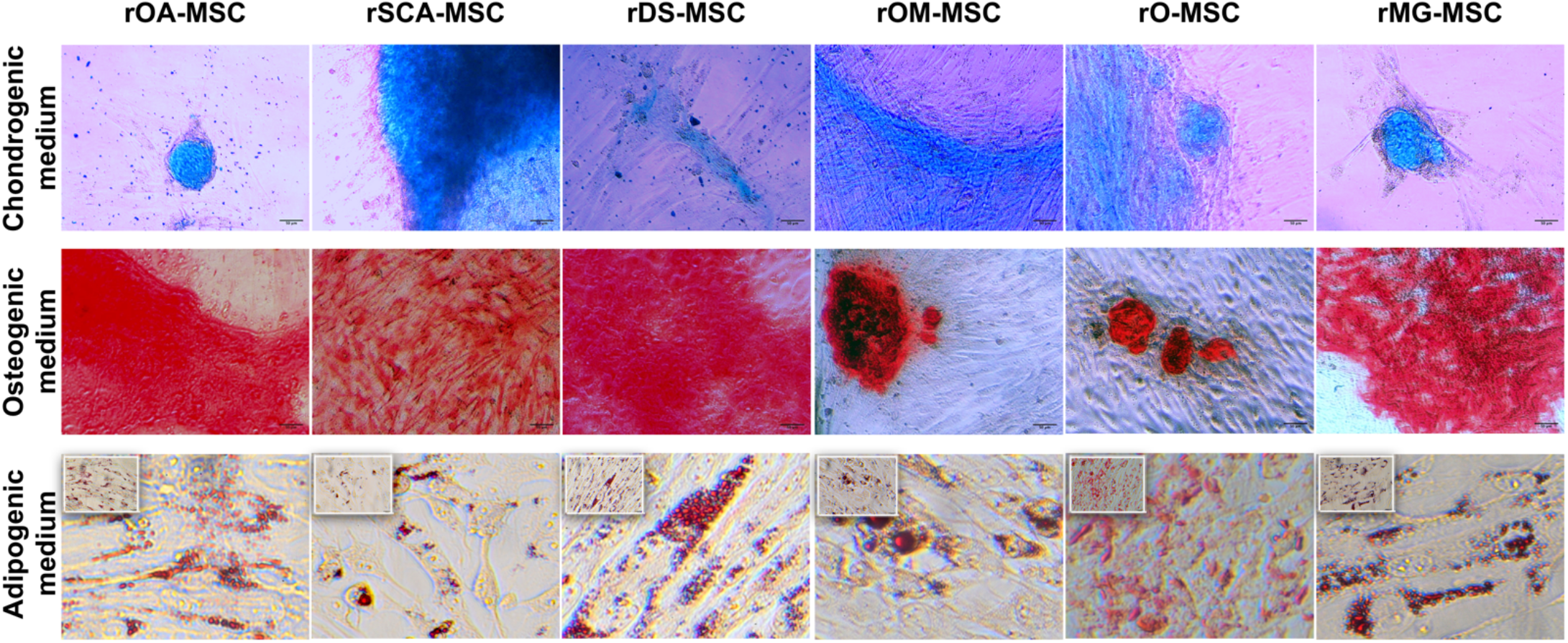
*In vitro* differentiation of rMSC to different lineages. Images show Alcian blue staining of acidic proteoglycan in cells cultured in chondrogenic differentiation medium (top panel); Alizarin Red S staining of calcium deposits in cells cultured in osteogenic differentiation medium (middle panels); and Oil red O staining of lipid droplets in cells cultured in adipogenic differentiation medium (bottom panels). Bright-field images were acquired with 200× magnification (bars = 50 µm) for top and middle panels and with 400x magnification (bars = 100 µm) for bottom panels.

## 3. Discussion

Our findings revealed that rabbit MSCs could be isolated and expanded *in vitro* from visceral adipose depots, subcutaneous fat, dermal skin, oral mucosa, oviduct, and mammary gland tissues of an adult female wild rabbit. Our results also demonstrate that rOA-MSCs, rSCA-MSCs, rDS-MSCs, rOM-MSC, rO-MSC, and rMG-MSCs shared similar characteristics in terms of morphology, expression of mesenchymal and pluripoten-cy-related markers, and differentiation ability into chondrocytes, osteocytes, and adipocytes. Passaged cells had a more homogeneous morphology and formed colonies as the culture progressed, compared to primary cultures. As we have shown in previous studies in porcine and bovine species, these morphological observations indicate that the isolated cells may contain both mature and progenitor populations [28–30].

Regarding cell surface markers characteristic of rMSC, contradictory results have been described in rabbits, as summarized by Zomer et al. [31]. Considering this, we have selected a panel of antibodies that have shown high sensitivity and repeatability and confirmed these results through RT-PCR assessment of expression at the mRNA level. Our data demonstrate that all rMSCs isolated were negative for CD34 and CD45 but positive for CD44, CD29, NANOG, OCT4, and SOX2, coinciding with the results reported in the literature for rMSCs isolated from different tissues: adipose [20], gingival [21], amniotic fluid [23] and bone marrow [25, 32].

The basic *in vitro* trilineage differentiation capacity of rMSC, that is adipocytes, osteocytes, and chondrocytes have been reported in the literature only for adipose [31] and for bone marrow rMSC [25, 26, 33]. Our results show that all isolated rMSC lines, regardless of their tissue of origin, were able to differentiate to the three mesodermal lineages, with lipid droplet accumulation, calcium deposits, or presence of proteoglycan in response to *in vitro* adipogenic, osteogenic, or chondrogenic stimuli, respectively.

The gene expression profile of adipose MSC from subcutaneous fat in Zucker rats was found to be distinct from the gene expression profiles of the other four visceral designated depots (epicardial, epididymal, mesenteric, and retroperitoneal), which clustered together but were not identical [34]. Also, differences between adipose-MSC from mesenteric and omental depots have been documented [35]. These findings show that individual visceral “subdepots” should not be considered interchangeable. Moreover, increased visceral (central) fat is more pathological, resulting in insulin resistance and chronic inflammation, whereas subcutaneous (peripheral) fat is more beneficial in systemic metabolism [36]. These differences are cell-intrinsic and persist in culture [35, 37], revealing the mechanisms underlying the pathophysiological differences between adipose depots. Individual adipose depots, which function as separate “mini-organs,” are formed and maintained by distinct A-MSC populations, with consistent differences in adipogenic potential and function later in life [38, 39]. We have however not observed major differences between rMSCs isolated from subcutaneous adipose tissue or ovarian adipose depot, although it would be interesting to make a more in-depth characterization.

Rabbit embryo and fetoplacental development are similar to human development, making the rabbit an excellent model for studying embryo-maternal communication, the effects of maternal metabolic disorders on offspring development and long-term health [40], and the development of *in vitro* models of pathogen infection during pregnancy. In 2009 Jazedje et al. showed for the first time that human fallopian tubes are a rich additional source of MSCs and these cells were designated as human tube MSCs (htMSCs) [41]. The presence of mesenchymal cell populations in this part of the female reproductive system has only been documented in humans until now [42–45]. Our rO-MSC will allow the development of both physiological and pathological *in vitro* models.

Due to the remarkable cyclical changes in proliferation, lactation, and involution that occur in the breast tissue throughout life and pregnancy, the field of mammary gland physiology has been historically interested in stem cell biology. Shackleton *et al*. demonstrated that a single cell lacking hematopoietic and endothelial antigens, and expressing high levels of CD29 could generate an entire mammary gland [46]. Following this description, in rMG-MSC we found no expression of the hematopoietic lineage markers CD34 and CD45, while they expressed CD29 and CD45 as well as pluripotency markers (NANOG, OCT4, and SOX2). Mammary gland stem cells have subsequently been shown to play key roles in both regeneration of the mammary gland and the development of mammary gland tumors. CD44 has been used as a marker of cancer-initiating cells in various cancers, including prostate, pancreas, and colon [47, 48]. Considering the above, rMG-MSC is a promising *in vitro* model for the study of both breast physiology and breast cancer, since it expressed CD44. Until now the presence of MSCs in the mammary gland has only been studied in mice [49–52].

The fact that rabbit hemorrhagic disease virus (RHDV), a relevant member of the Caliciviridae family, cannot be propagated *in vitro*, has greatly hampered the progress of investigations into its mechanisms of pathogenesis, translation, and replication. In this regard, the isolation of new rMSC lines from target tissues for the main viral pathogens of the species (myxoma virus and RHDV) will allow an in-depth analysis of both their susceptibility to viral infection and their possible role in the regulation of host antiviral response.

## 4. Materials and Methods

### 4.1. Isolation and culture of rMSCs

Different tissue samples were obtained postmortem from an adult female wild rabbit (*Oryctolagus cuniculus*).

For the isolation of primary mesenchymal cultures from oral mucosa (rOM-MSC), dermal skin tissue (rDS-MSC), subcutaneous adipose tissue (rSCA-MSC), ovarian adipose tissue (rOA-MSC), oviduct (rO-MSC), and mammary gland (rMG-MSC), the procedures are detailed in Calle et al. [28]. Briefly, the collected samples were minced before the tissues were incubated in a collagenase type I solution. Thereafter, a volume of culture medium was added to block the action of collagenase. The resulting pellets were resuspended in culture medium and plated in a 100-mm^2^ tissue culture dish (Jet-Biofil, Guangzhou, China) and incubated in an atmosphere of humidified air and 5% CO2 at 37 °C. The culture medium was changed every 48–72 h. Isolated colonies of putative rMSCs were apparent after 5-6 days in culture and were maintained in growth medium until ∼ 75% confluence. The cells were then treated with 0.05% trypsin–EDTA (T/E) and further cultured for subsequent passage in 100-mm^2^ dishes at 50,000 cells/cm^2^.

### 4.2. Immunocytochemical analysis by flow cytometry

Cell cultures at 80–90% confluence were detached and fixed with 4% paraformaldehyde for 10 min. Stainings were performed as detailed in Calle et al. [28], employing anti-CD44 (clone W4/86, Bio-Rad); anti-CD34 (Mouse monoclonal, Invitrogen), and anti-CD45 (clone L12/201, Bio-Rad). Appropriated secondary staining using F(ab’)2-Goat anti-Mouse IgG (H+L) Cross-Adsorbed Secondary Antibody, APC (A10539) from Life Technologies. Negative control samples corresponded to samples in which the primary antibody was omitted. Samples were analyzed in a FACSCalibur (BD Biosciences) using Flow-JoX Software® version 10.0.7r2.

### 4.3. RT-PCR analysis of characteristic mesenchymal mRNA expression in rMSC

Total RNA was extracted from 2,5 ·10^6^ cells by RNeasy Mini kit (Qiagen), according to the manufacturer’s instructions. The RNA starting template was analyzed for integrity and quantity by Nanodrop test, and 600 ng were used in the RT-PCR reaction. cDNA was synthesized and amplified using OneStep RT-PCR Kit (Qiagen), according to the manufacturer’s instructions, and using the following gene-specific primers: Rabbit CD29, forward (AGAATGTCACCAACCGTAGCA), reverse (CACAAAGGAGCCAAACCCA); rabbit CD44, forward (TCATCCTGG-CATCCCTCTTG), reverse (CCGTTGCCATTGTTGATCAC); rabbit CD45, forward (TACTCTGCCTCCCGTTG), reverse (GCTGAGTGTCTGCGTGTC); rabbit CD34, forward (TTTCCTCATGAACCGTCGCA), reverse (CGTGTTGTCTTGCGGAATGG); NANOG, forward (GCCAGTCGTGGAGTAACCAT), reverse (CTGCATGGAGGACTGTAGCA); OCT4, forward (GAGGAGTCCCAGGACATGAA), reverse (GTGGTTTGGCTGAACACCTT); SOX2, forward (CAGCTCGCAGACCTACATGA), reverse (TGGAGTGGGAGGAAGAGGTA) [23]. PCR products were separated on a 1.5% agarose gel (MB Agarose, Bio-Rad) stained with GelRed® Nucleic Acid Gel Stain (Biotium, #41002).

### 4.4. In vitro differentiation potential assay

For adipogenic, osteogenic, and chondrogenic differentiation, cells were cultured according to the manufacturer’s instructions of the StemPro® adipogenesis, osteogenesis, or chondrogenesis differentiation kits, respectively (Thermo Fisher Scientific, Meridian Road Rockford, IL) and analyzed as detailed in Calle et al. [28].

## 5. Conclusions

In summary, this study describes the isolation and characterization for the first time of cell lines from different tissue origins with a clear mesenchymal pattern, derived from a natural rabbit population within the species native region. We show that rMSCs do not present differences in terms of morphological features, expression of the cell surface, and intracellular markers of pluripotency and *in vitro* chondrogenic, osteogenic and adipogenic differentiation capacities attributable to their tissue of origin.

## Author Contributions

Conceptualization, M.Á.R.; methodology, M.Á.R., E.B. and A.C.; software, M.Á.R. and A.C.; validation, M.Á.R., E.B., J.B., and A.C.; formal analysis, M.Á.R. E.B. and A.C.; investigation, M.Á.R., E.B., J.B., M.Z. and A.C.; resources, M.Á.R., E.B., J.B., M.Z. and A.C; data curation, M.Á.R. E.B. and A.C.; writing—original draft preparation, M.Á.R.; writing—review and editing, M.Á.R., E.B., J.B., M.Z. and A.C; visualization, M.Á.R., and A.C.; supervision, M.Á.R.; project administration M.Á.R., E.B. and J.B.; funding acquisition, M.Á.R., E.B. and J.B. All authors have read and agreed to the published version of the manuscript.

## Acknowledgments

The authors are grateful to Dra María Yáñez-Mó (Dpto. de Biología Molecular, UAM, Madrid, Spain) for critical reading of the manuscript.

The authors thank Marta Díaz-de Frutos for her experimental assistance during her stay at CISA, INIA.

